# Vitamin D is Glucoprotective in Aging Males but Not Females

**DOI:** 10.1101/2024.02.13.580162

**Authors:** Olivia Z.B. Ginnard, Maria Morales, Ji Youn Youn, Yong Xu, Stephanie R. Sisley

## Abstract

Type 2 diabetes is strongly linked to vitamin D deficiency in older adults. However, there is a discrepancy between clinical trials in adults on the efficacy of vitamin D treatment in prediabetes and diabetes. In addition, human data indicates there may be sexual dimorphism in the effect of vitamin D deficiency on dysglycemia that is more pronounced in men. These incongruities may be due to our limited understanding of the underlying mechanisms of vitamin D in glucose homeostasis among its vast target tissues across the body. Furthermore, vitamin D deficiency is diagnosed by low levels of a storage form of vitamin D, which may not be an accurate indicator of vitamin D status in these individuals. Thus, measuring expression levels of vitamin D receptor (VDR)-target genes across tissues involved in glucose regulation and vitamin D pathways may be a more promising marker of vitamin D status. Here we describe the sex-specific physiological effects of vitamin D supplementation in an aged, non-obese mouse model on glucose homeostasis and tissue-specific gene regulation.

## INTRODUCTION

Vitamin D is a fundamental component of many basic physiologic processes vitally important to human health. Unsurprisingly, vitamin D deficiency is linked to poorer health outcomes in older adults. Numerous studies show a negative correlation between vitamin D levels and the development of type 2 diabetes (T2D) [1-6]. T2D is one of the most prevalent chronic diseases in adults in the U.S. over the age of 65 with an increase in prevalence by 62% over the last decade resulting in a staggering $104 billion per year in medical costs [7]. T2D in the geriatric population is associated with higher mortality and morbidity likely due in-part to complications of polypharmacy in an aging and a more cognitively impaired population [7]. Given the link between vitamin D deficiency and T2D, dietary vitamin D may be a safe and cost-effective therapy in the prevention and treatment of T2D in the aging adult population. Interestingly, a recent meta-analysis confirms that vitamin D can prevent the progression of prediabetes in lean individuals [8].

Through ingestion or skin absorption, vitamin D is introduced to the body primarily as cholecalciferol and then hydrolyzed to the storage form, 25-hydroxyvitamin D, in the liver. It is further hydrolyzed to the most biologically active form of 1,25-dihydroxyvitmain D (1,25D) in the kidney. 1,25D then binds to the vitamin D receptor (VDR) to induce changes in the transcription of hundreds of genes [9-24]. Current literature suggests that the VDR has an integral role in multiple tissues. This includes prevention of inflammation in beta-cells and the liver, glucose tolerance regulation in the brain, and restoration of insulin signaling from obesity-induced derangements in skeletal muscle [25-28].

Despite these findings, there is a discrepancy between clinical trials in adults on the efficacy of vitamin D treatment in prediabetes and diabetes [29-31]. This may be due to our limited understanding of the underlying mechanistic differences in tissue-specific vitamin D regulation, specifically the role of vitamin D-regulated gene transcription in glucose homeostasis across multiple tissues and the effect of hyperglycemia on beneficial effects of vitamin D action.

Here, we aim to describe the physiological effects of vitamin D supplementation in an aged, non-obese mouse model on glucose homeostasis and tissue-specific gene regulation.

## MATERIALS AND METHODS

### Animals

Mice were group-housed at Baylor College of Medicine (BCM) on a 12 h light/dark cycle with ad libitum access to food and water. The mice used for these studies were on a C57BI/6J background. All studies used littermate controls. For an aged phenotype, mice were 37-47 weeks (middle-aged mice) at the start of the diet supplementation (see below) and 60-70 weeks at the end of the experiment. Mice were euthanized with intraperitoneal sodium pentobarbital per BCM protocols. All studies were approved by the BCM Institutional Animal Care and Use Committee (IACUC).

### Diet

Male and female mice were studied in a lean state utilizing a chow diet background (15 kcal%fat, Research Diets D11112201). The animals were further stratified to three different dietary vitamin D intakes (within the chow background): low (100 IU/kg diet), normal (1000 IU/kg diet), and high (5000 IU/kg diet). The doses of vitamin D were based on literature review of mouse models utilizing various vitamin D concentrations to achieve dose-dependent physiologic changes in vitamin D but prevent hypo/hypercalcemia from occurring [32-35]. The vitamin D doses were also chosen to mimic standard treatment doses in humans for vitamin D deficiency without causing vitamin D toxicity. The mice were on the stratified vitamin D diets for 7 weeks prior to glucose tolerance testing [Fig 1].

**FIGURE 1:**
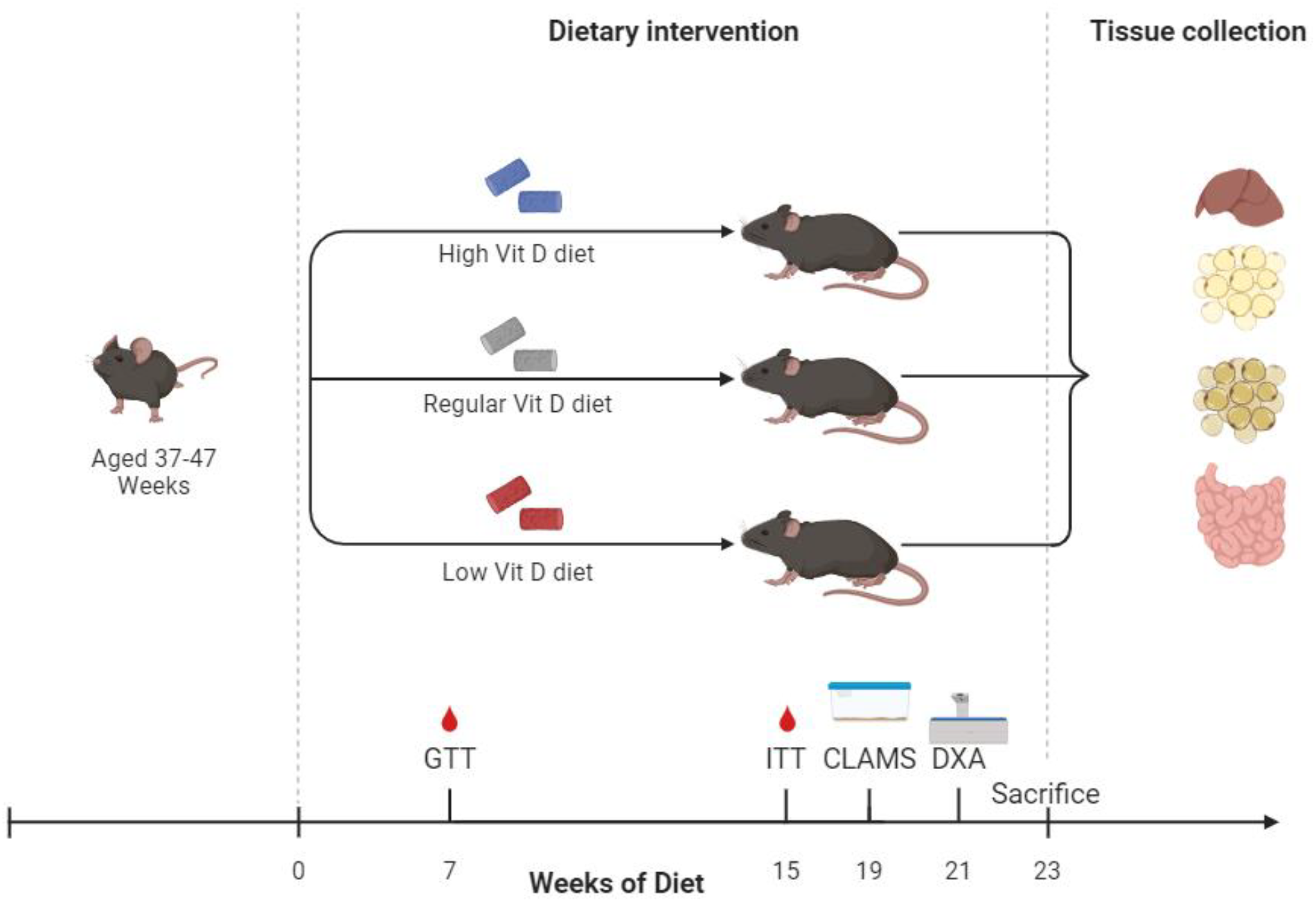
Research Design Schematic (*image created with BioRender*.*com*)

### Glucose and Insulin Tolerance Tests

For glucose tolerance testing (GTT), mice were fasted for four hours to empty the stomach. They then received 1.0 g/kg dextrose (D20W) intraperitoneal injection. Blood glucose from a tail laceration was measured in duplicate at time 0 (prior to dextrose administration), 15, 30, 45, 60, and 120 minutes after dextrose administration via glucometer. Eight weeks after glucose tolerance testing, mice underwent insulin tolerance testing (ITT). A similar experimental set-up as the glucose tolerance test was utilized with exception of 1.0 unit/kg Humulin-R insulin given instead of dextrose, and blood glucose was measured prior to and at 15, 30, 45, and 60 minutes after insulin administration. Both tests were performed in freely moving, conscious mice.

### Body Composition and Energy Expenditure

Four weeks after insulin tolerance testing, mice underwent energy expenditure testing utilizing the Comprehensive Laboratory Animal Monitoring System (CLAMS) cages. Animal underwent acclimation in the CLAMS cages for two days before obtaining energy expenditure data. After 21 weeks of stratified vitamin D diet, we indirectly measured bone mineral content via PIXImus.

### Gene Expression

At time of euthanasia, subcutaneous fat (SAT), visceral fat (VAT), liver, and the first 5 cm of small bowel tissue (SBT) were grossly dissected and rapidly frozen with liquid nitrogen. Tissue RNA was extracted using a Qiagen RNeasy kit. cDNA was isolated and real-time quantitative PCR (qPCR) was performed using a TaqMan 7900 sequence detection system with TaqMan universal PCR master mix and TaqMan gene expression assays (Applied Biosystems). Relative gene expression for *Vdr, Pparg, Cyp27b1, Trpv6, S100g, G6p, Gck, Insr*, and *Glut4* genes was calculated relative to the housekeeping gene *Gapdh* using the ΔΔCT method [36].

### Statistics

For GTT, ITT, body weight curves, and energy expenditure, data were analyzed with a paired 2-way ANOVA. For AUC, data were analyzed with one-way ANOVA. The data for qPCR measurements were analyzed utilizing simple linear regression analysis. P < 0.05 was considered to be statistically significant.

## RESULTS

### Dietary Vitamin D Did Not Significantly Impact Body Weight, Bone Mineral Content, or Energy Expenditure

There were no significant differences in body weight [Fig. 2a-b] between the different vitamin D diet cohorts in either males or females. Despite this, we found a significant increase in the food intake of the high dose vitamin D diet after ten weeks of stratified diets [Fig 2c-d]. This was not accounted for by a change in energy expenditure as we did not observe any differences between the groups in respiratory exchange ratio (RER), energy expenditure, or total activity [Supplemental Fig 1a-f]. Furthermore, bone mineral content was similar across dose cohorts [Fig 2e-f].

**FIGURE 2:**
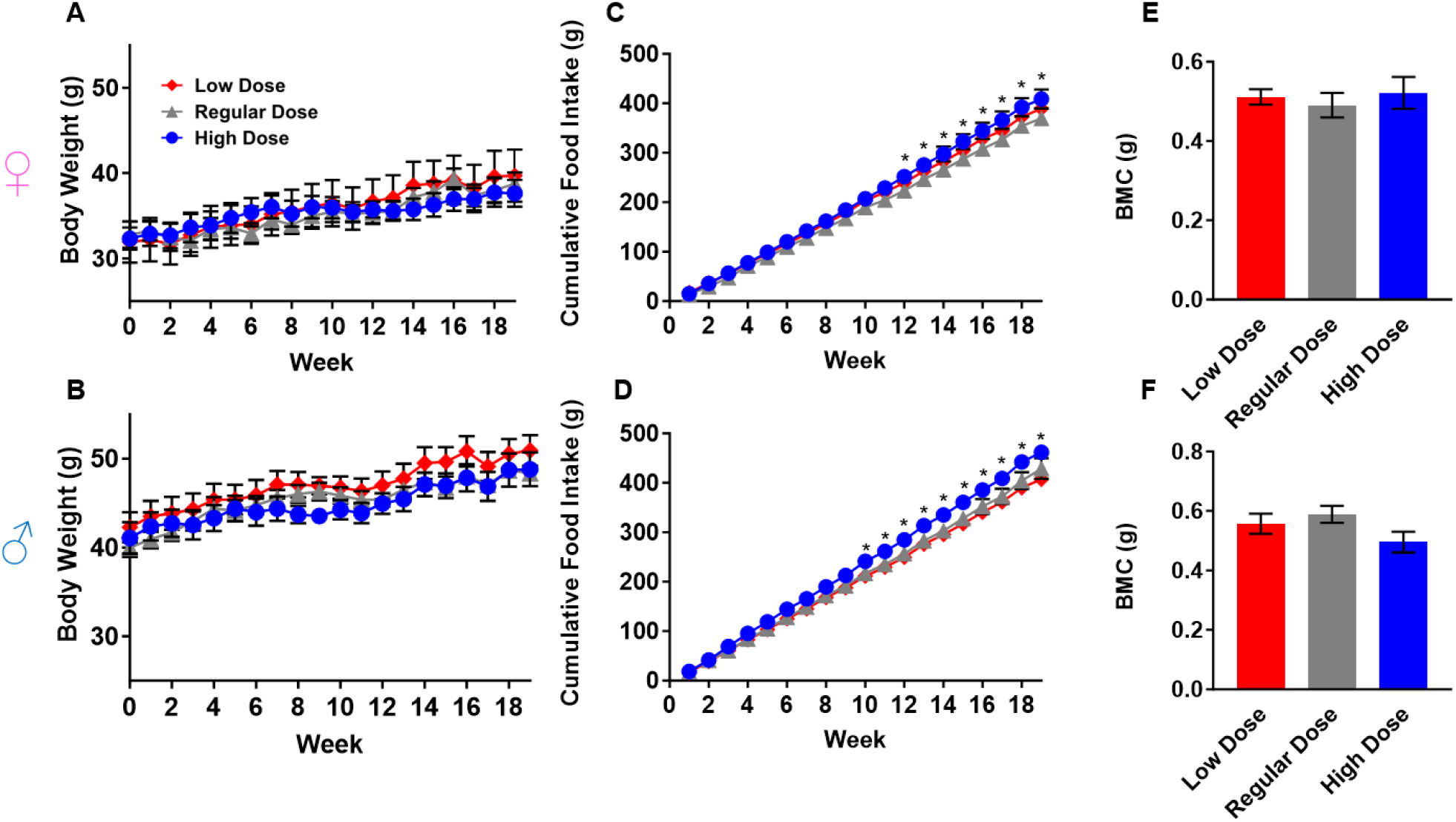
Different dietary vitamin D supplementation did not affect anthropometric measurements. **(A)** Body weights in lean female mice and **(B)** male mice on low (100 IU/kg), normal (1000 IU/kg), and high (5000 IU/kg) dietary vitamin D supplementation. **(C)** Cumulative food intake of lean female mice and **(D)** male mice on the stratified dietary vitamin D supplementations. **(E)** Bone mineral content via DXA measurements of both lean female mice and **(F)** male mice on the stratified vitamin D diets. \**p*<0.05.

### High Dose Dietary Vitamin D Impacts Glucose Tolerance and Insulin Tolerance

Despite having no difference in body weight, male mice consuming a high dose vitamin D diet had improved glucose tolerance as compared to mice on low or regular dose vitamin D diets [Fig 3a-b]. Similarly, the male mice eating a high dose vitamin D diet had better insulin sensitivity as compared to mice on the low dose vitamin D diet [Fig 3c-d]. Female mice did not show any significant differences in glucose tolerance or insulin sensitivity when stratified by diet [Fig 3e-h].

**FIGURE 3:**
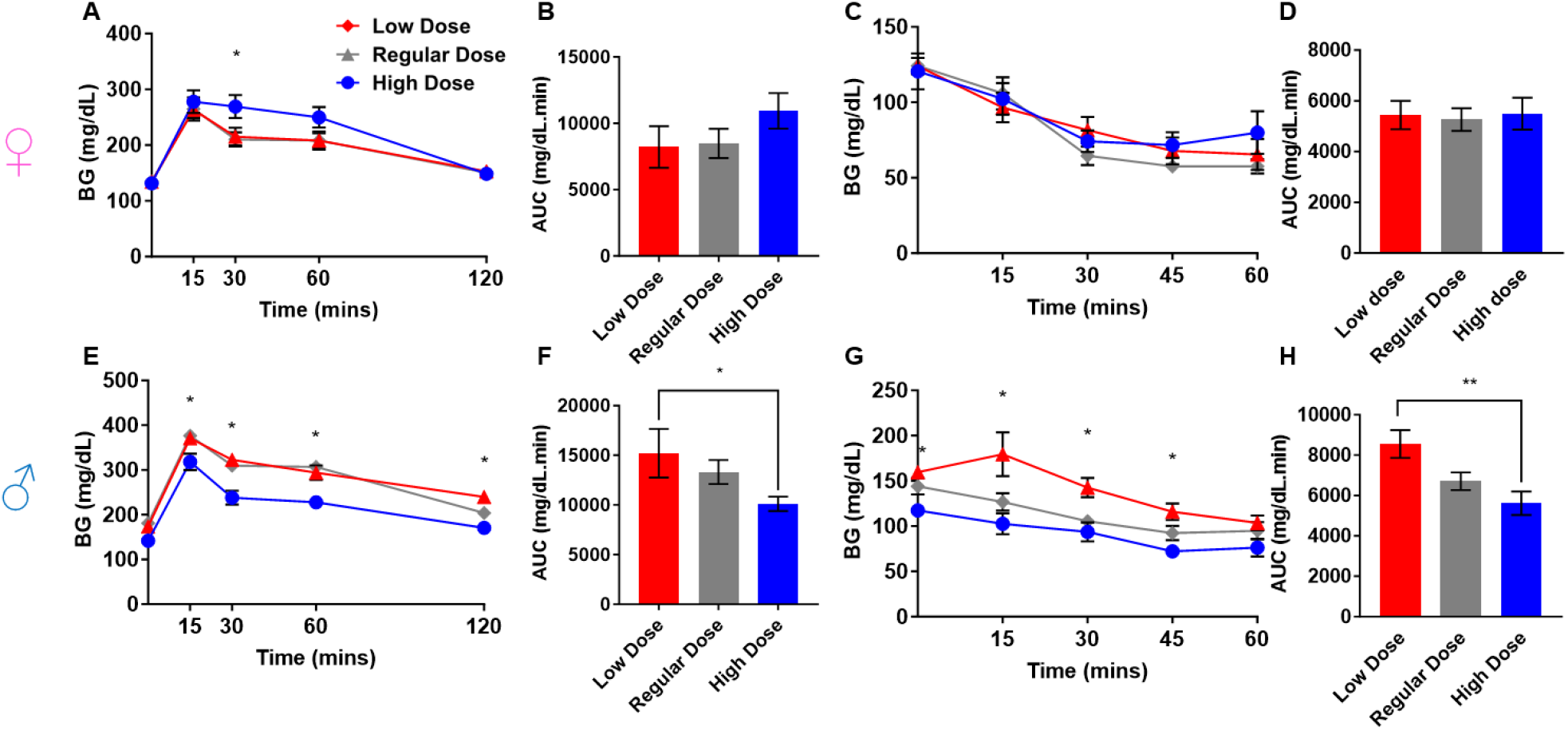
Dietary vitamin D impacts glucose tolerance and insulin tolerance. **(A)** Glucose tolerance test (1.0 g/kg) with **(B)** area under the curve in 45-55 week old lean female mice. **(C)** Insulin tolerance test (1.0 unit/kg) with **(D)** area under the curve in 53-63 week old lean female mice. **(E-H)** Same glucose and insulin testing in age-matched lean male mice. \**p*<0.05; ** *p*<0.01.

### VDR Gene Expression Correlates to Glucose Metabolism-Related Gene Expression

To determine the role of vitamin D in glucose tolerance at a molecular level, we evaluated the effects of diet on gene expression. Since gene regulation is tissue specific, we analyzed the expression of *Vdr* in all tissues but chose VDR-related tissue-specific genes for each tissue. These included *Trpv6, Cyp27b1, and S100g* in the SBT and *Pparg* in the adipose tissues [13-14 19, 37-40]. In addition, we analyzed expression of genes crucial to glucose and insulin regulation in the body, such as *Insr* and *Glut4* in all tissues and additionally *Gck* and *G6p* in liver tissue [41-42]. Tissue-specific and glucoregulatory gene expressions were compared to *Vdr* expression within each tissue to appreciate the changes in expression levels of these tissue-specific or gluco-regulatory-specific genes in relation to genomic actions of *Vdr*, as influenced by *Vdr* levels.

In SAT, we found significant correlation in gene expression levels of *Vdr* with both *Insr* and *Glut4* [Supplementary Fig 2a-d]. We did not find any significant gene expression differences when stratified by vitamin D dose diet [Supplementary Fig 2e]. Interestingly in the VAT, we again found that *Vdr* gene expression levels correlated with *Insr* gene expression but additionally with *Cyp27b1* gene expression [Supplementary Fig 2f-i], a key gene responsible for the final step of producing calcitriol, the most biologically active endogenous VDR ligand. When further stratified by diet, we found that gene expression levels of *VDR* were significantly higher in the cohort on the low dose vitamin D diet [Supplementary Fig 2j].

In SBT, we found significant correlation between gene expression levels of *Vdr* and *Insr, Glut4*, and additionally *S100g* [Supplementary Fig 3a-e]. However, we did not find any significant gene expression differences in glucose-related genes in SBT when stratified by vitamin D dose diet [Supplementary Fig 3k]. Finally, in liver tissue, we again found the *Vdr* gene expression levels correlated with *Insr* and *Glut4* gene expression [Supplementary Fig 3f-j] without any significant gene expression differences when stratified by vitamin D dose [Supplementary Fig 3l].

### Sexual Dimorphism in VDR Role in Glucose Metabolism

To investigate whether the observed sex differences in glucose tolerance on different vitamin D diets are explained by response in gene transcription, we performed similar analyses of gene expression within each individual sex. In SAT of both sexes, *Vdr* expression significantly correlated with both *Insr* and *Pparg* [Fig 4a-d, f-i]. In females, *Vdr* expression also significantly correlated with *Glut4*. When further stratified by diet, mice on low vitamin D diet had lower levels of *Pparg, Vdr*, and *Insr* expression compared to regular vitamin D diet in males [Fig 4e]. Females had no significant gene differences when stratified by diet [Fig 4j]. In VAT tissue, the only correlation with *Vdr* gene expression occurred with *Cyp27b1* in females [Fig 5g]. No other significant gene expression correlations found between the sexes [Fig 5a-d, f-i]. However, when stratified by dietary vitamin D dose, males, but not females, on low dose vitamin D diet had lower gene expression levels of *Glut4* [Fig 5e, j].

**FIGURE 4:**
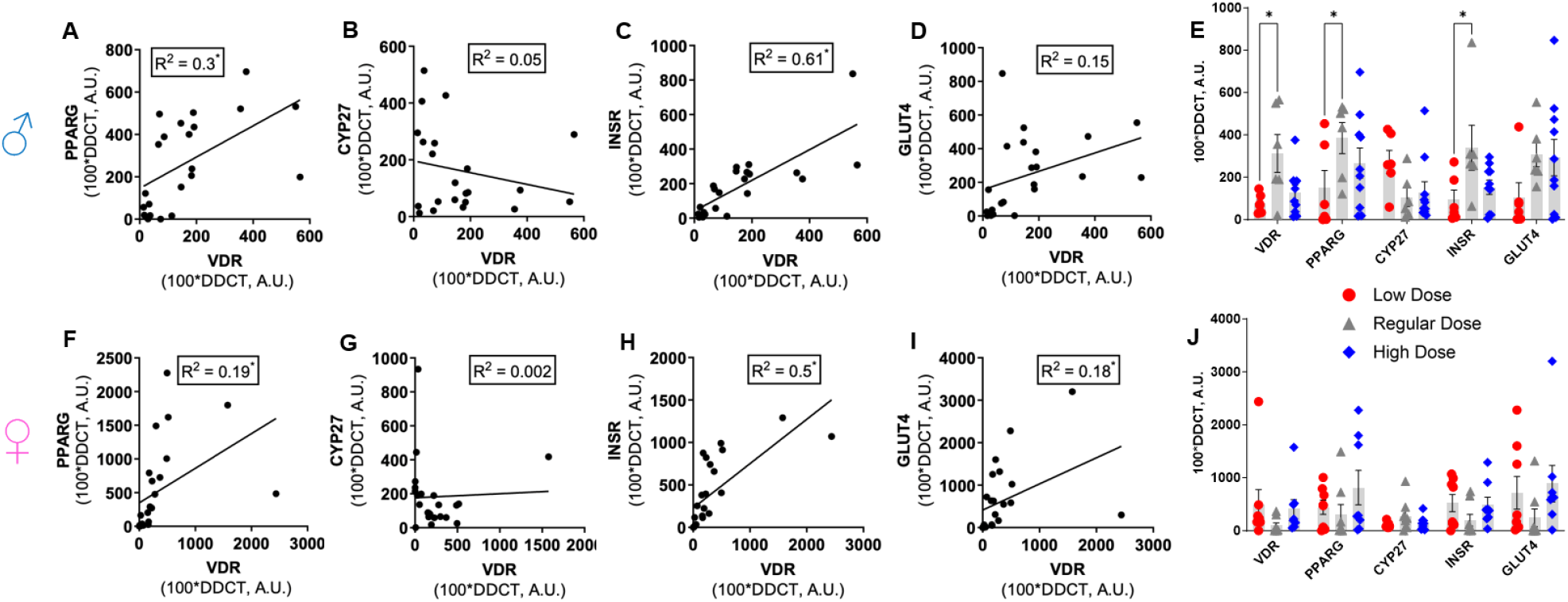
Sex differences in VDR Gene expression as correlated to glucose metabolism-related gene expression in SAT. **(A-D)** Male SAT gene expression of *Vdr* vs *Pparg, Cyp27a1, Insr*, and *Glut4* gene expression, respectively. **(E)** All examined SAT genes in males stratified by diet. **(F-I)** Female SAT gene expression of *Vdr* vs *Pparg, Cyp27a1, Insr*, and *Glut4* gene expression, respectively. **(J)** All examined SAT genes in females stratified by diet. All expression values calculated relative to the housekeeping genes *Gapdh* using the ΔΔCT method. \**p*<0.05.

**FIGURE 5:**
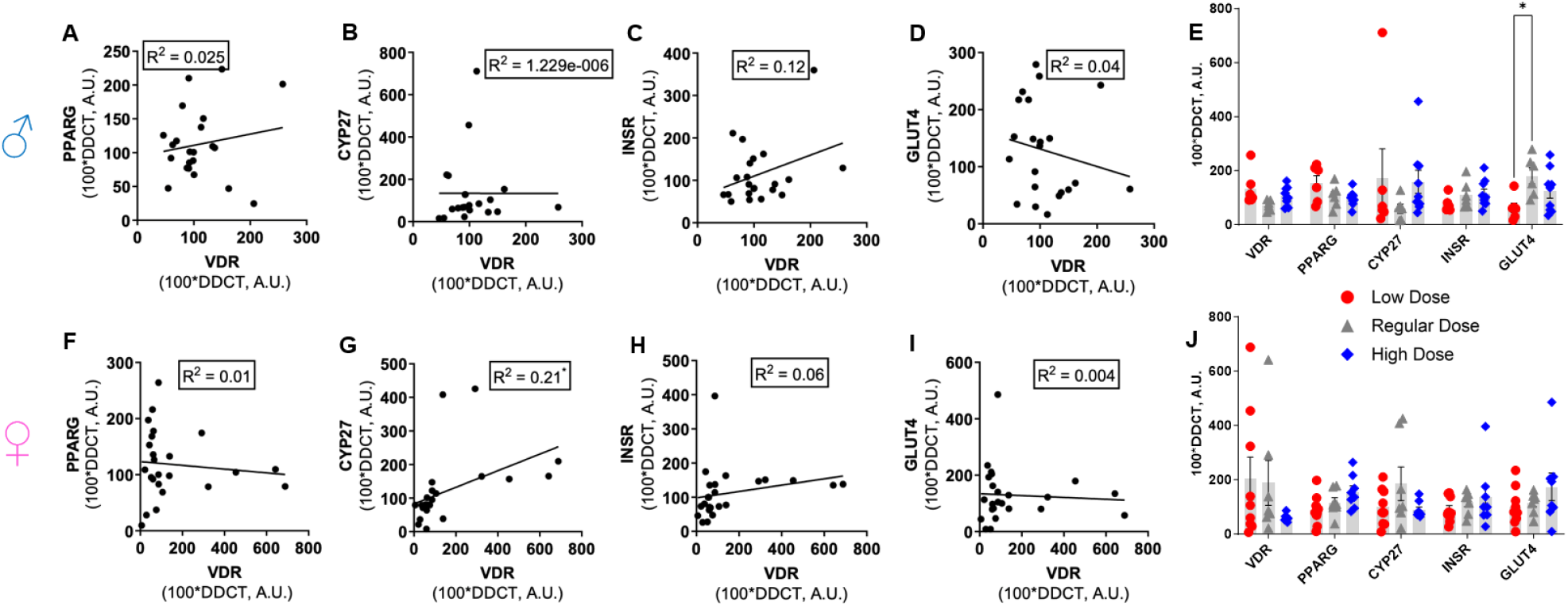
Sex differences in VDR Gene expression as correlated to glucose metabolism-related gene expression in VAT. **(A-D)** Male VAT gene expression of *Vdr* vs *Pparg, Cyp27a1, Insr*, and *Glut4* gene expression, respectively. **(E)** All examined VAT genes in males stratified by diet. **(F-I)** Female VAT gene expression of *Vdr* vs *Pparg, Cyp27a1, Insr*, and *Glut4* gene expression, respectively. **(J)** All examined VAT genes in females stratified by diet. All expression values calculated relative to the housekeeping genes *Gapdh* using the ΔΔCT method. \**p*<0.05.

In the SBT of both sexes, we again found that *Vdr* gene expression correlated with *Insr* gene expression [Fig 6d,j]. Additionally, male mice had a significant correlation between levels of *Vdr* gene expression and *Trpv6*, a gene crucial to the absorption of vitamin D in the intestine, and female mice had significantly correlated gene expression levels of *Vdr* and *Glut4* [Fig 6a-e, g-k]. While *Vdr* and *Insr* in both sexes and *S100g* in male mice trended towards significant upregulation in mice on a high vitamin D diet, neither sex had any significant gene expression differences in glucose-related genes when stratified by diet [Fig 6f,l]. Finally, in the liver, significant correlations of *Vdr* gene expression occurred with *Insr* and *G6p* expression levels in male but not female mice [Fig 7a-e, g-k]. When stratifying by vitamin D dose diet, male mice on the high vitamin D dose diet had significantly higher gene expression levels of *Insr* compared to low dose and regular dose vitamin D diet [Fig 7f]. No significant differences in gene expression levels in the liver were observed in female mice when stratified by diet [Fig 7l].

**FIGURE 6:**
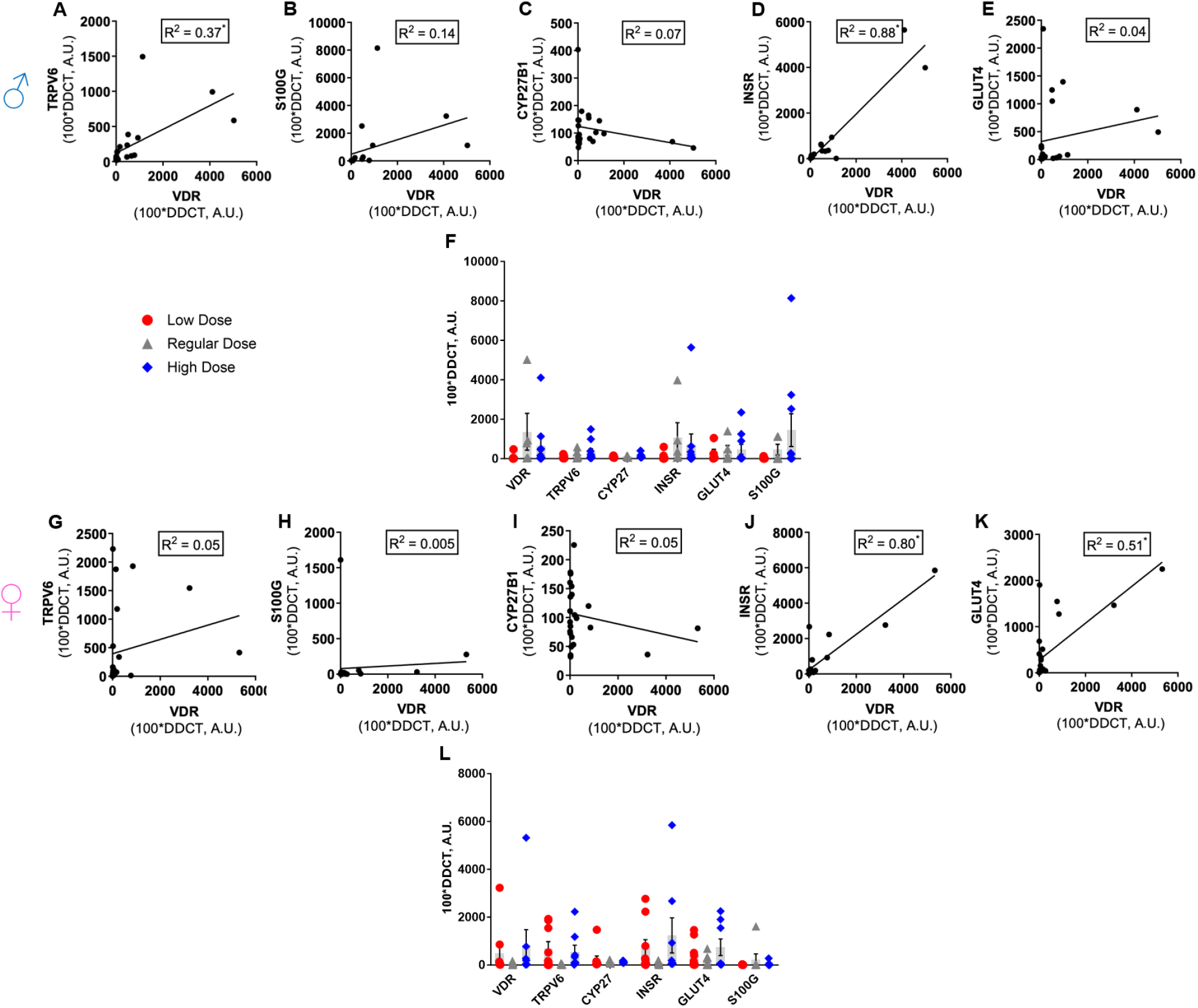
Sex differences in VDR Gene expression as correlated to glucose metabolism-related gene expression in SBT. **(A-E)** Male SBT gene expression of *Vdr* vs *Trpv6, S100g, Cyp27b1, Insr*, and *Glut4* gene expression, respectively. **(F)** All examined SBT genes in males stratified by diet. **(G-K)** Female SBT\ gene expression of *Vdr* vs *Trpv6, S100g, Cyp27b1, Insr*, and *Glut4* gene expression, respectively. **(L)** All examined SBT genes in females stratified by diet. All expression values calculated relative to the housekeeping genes *Gapdh* using the ΔΔCT method. \**p*<0.05.

**FIGURE 7:**
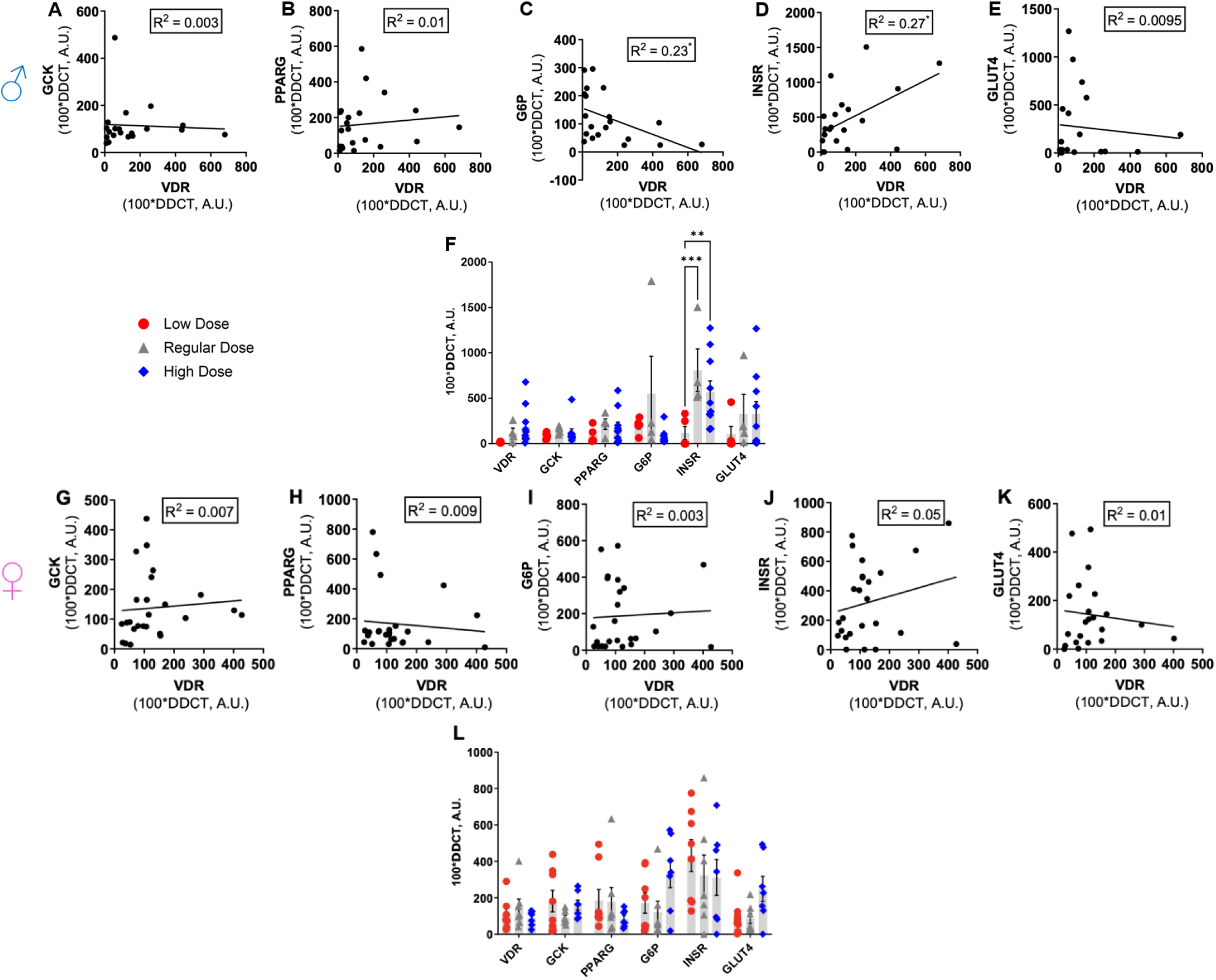
Sex differences in VDR Gene expression as correlated to glucose metabolism-related gene expression in liver tissue. **(A-E)** Male liver gene expression of *Vdr* vs *Gck, Pparg, G6p, Insr*, and *Glut4* gene expression, respectively. **(F)** All examined liver genes in males stratified by diet. **(G-K)** Female liver gene expression of *Vdr* vs *Gck, Pparg, G6p, Insr*, and *Glut4* gene expression, respectively. **(L)** All examined liver genes in females stratified by diet. All expression values calculated relative to the housekeeping genes *Gapdh* using the ΔΔCT method. \**p*<0.05; ** *p*<0.01; *** *p*<0.001.

## DISCUSSION

Our results show that dietary vitamin D supplementation can improve glucose tolerance and insulin sensitivity in lean, aged male mice. These findings replicate a previous meta-analysis of lean adults indicating the benefit of vitamin D in preventing prediabetes. While there is little data on sex-specific effects of vitamin D in humans (discussed more below), we also show here that dietary vitamin D has sex-specific effects in mice, and this may be due to effects of vitamin D on tissue-specific gene expression. To our knowledge, this study is the first to describe a glucose phenotype in a lean and aged mouse model on various dietary vitamin D doses. This is significant, as it may provide primary care providers a safe and cost-effective option to offer aging adult patients at risk of prediabetes or T2D, which is associated with a higher risk of mortality and morbidity than their younger counterparts.

Changes in dietary vitamin D did not change body weight across cohorts. This indicated that the glucose effects were primarily due to direct effects of dietary vitamin D and not to obesity or insulin resistance related to fat accumulation changes. While we did not see differences in bone mineral content across the treatment groups, this is expected given that bone mineral content changes would likely require more than 12 weeks of treatment [43]. Due to issues with sample storage during the pandemic, we were not able to determine 25-OHD and calcium levels from the serum of the various dietary vitamin D cohorts. However, given that body weights and bone mineral content were similar across groups, it is unlikely we missed significant hypo- or hypercalcemia.

Human data shows conflicting reports regarding vitamin D supplementation on glucose regulation; however there have been few mechanistic studies to investigate this [8, 29-31]. Human studies have shown that high dose vitamin D supplementation can change gene expression levels of VDR target genes in SAT and peripheral blood [11, 17, 22]. Furthermore, it has been shown that vitamin D is involved in the regulation of insulin synthesis and secretion [25-28, 44]. However, there have been no studies investigating the effects of dietary vitamin D at physiological ranges on gene expression across a variety of tissues nor in the context of glucose homeostasis. We investigated whether changes in dietary vitamin D differentially impacted gene expression in SAT, VAT, SBT and liver. Gene expression of *Insr* correlated with *Vdr* gene expression and was upregulated with dietary vitamin D across many tissues in the body, indicating that vitamin D supplementation might improve insulin action at these various tissues. Additionally, we found that *Glut4* gene expression correlated with *Vdr* gene expression in subcutaneous adipose tissue, in the small bowel, and in the liver. Since *Glut4* increases glucose uptake into cells, this suggests that dietary vitamin D may help prevent diabetes by improving glucose transport. These studies are the first in our knowledge to investigate VDR-target gene expression systematically, across multiple tissues, and in relation to glucose metabolism-related gene expression. These findings further clarify the mechanistic relationship between vitamin D deficiency and poorer glycemic control. A key outstanding question is whether these effects of dietary vitamin D are lost in the presence of existing diabetes or in an insulin resistant state, such as obesity.

Interestingly, the glucoprotective effects of dietary vitamin D supplementation were seen more prominently in male mice. Although both sexes had gene expression of *Vdr* correlate to expression of glucose metabolism-related genes of *Insr* and *Glut4* across multiple tissues, we saw that in male mice, *Glut*4 and *Insr* were also upregulated with dietary vitamin D supplementation in VAT and in the SAT and the liver, respectively. This indicates there may be sexual dimorphism in the effect of vitamin D supplementation on insulin action and glucose transport. Specifically, it appears that dietary vitamin D may be a key mediator in upregulating glucose homeostasis pathways in males. This finding is congruent to the limited literature exploring sex differences in vitamin D effects. Specifically, human data shows that vitamin D deficiency is linked to dysglycemia that is more pronounced in men [45-46]. In addition, Sisley et al. discovered that centrally delivered vitamin D improved glucose tolerance in male mice [47]. Other external factors are unlikely to have influenced the dietary differences in glucose and insulin tolerance in our study as animals were littermates, age-matched, and exposed to the same housing conditions.

Limitations of our studies include lack of blood samples to determine 25-OHD and calcium levels of the various dietary vitamin D cohorts. As these studies were conducted during the pandemic, we encountered unique issues such as appropriate access and management of blood sample storage. Thus, analyses were severely limited due to sample destruction.

In conclusion, this study shows that dietary vitamin D supplementation is crucial in preventing glucose intolerance and insulin resistance in lean, aged male mice. Further work should investigate the effect of hyperglycemia and insulin resistance on vitamin D action in response to glucose control and VDR-target gene expression across multiple tissues of the body. This study may provide the first step in utilizing vitamin D more effectively in the population that most desperately needs it – older adults with T2D.

## ACKNOWLEDGEMENTS

We thank Dr. Marta Fiorotto, director of the Mouse Metabolic Research Unit at the Children’s Nutrition Research Center, for her expertise and technical assistance with the Comprehensive Laboratory Animal Monitoring System (CLAMS) to measure metabolic physiology and the PIXImus to indirectly measure bone mineral content. We thank Terri Semien along with the rest of the Center for Comparative Medicine Staff for their excellent technical assistance with breeding and maintenance of mouse colonies used in this study.

## AUTHOR CONTRIBUTIONS

All authors listed have made a substantial, direct, and intellectual contribution to the work, and approved it for publication.

