## Supplemental Figures for "Vitamin D is Glucoprotective in Aging Males but Not Females"

Supplemental Materials

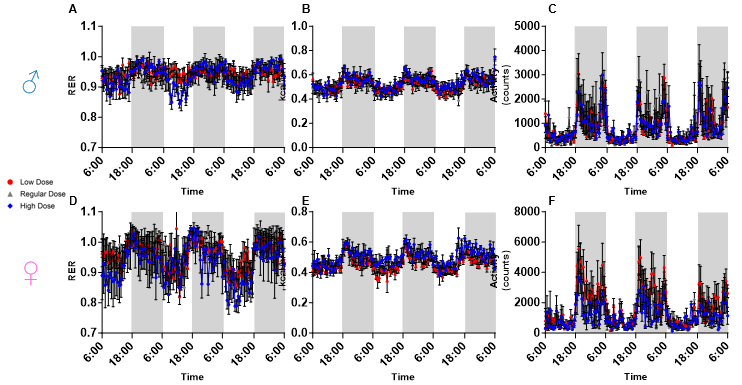

**SUPPLEMENTARY FIGURE 1**: Different dietary vitamin D supplementation did not affect energy expenditure measurements. **(A)** Respiratory Exchange Ratio (RER), **(B)** Energy Expenditure (Kcal/hr), and **(C)** total activity (counts) in lean male mice had no significant differences when compared by low (100 IU/kg), normal (1000 IU/kg), and high (5000 IU/kg) dietary vitamin D supplementation. Similar pattern was seen in the female mice **(D-F)**. **p*<0.05.

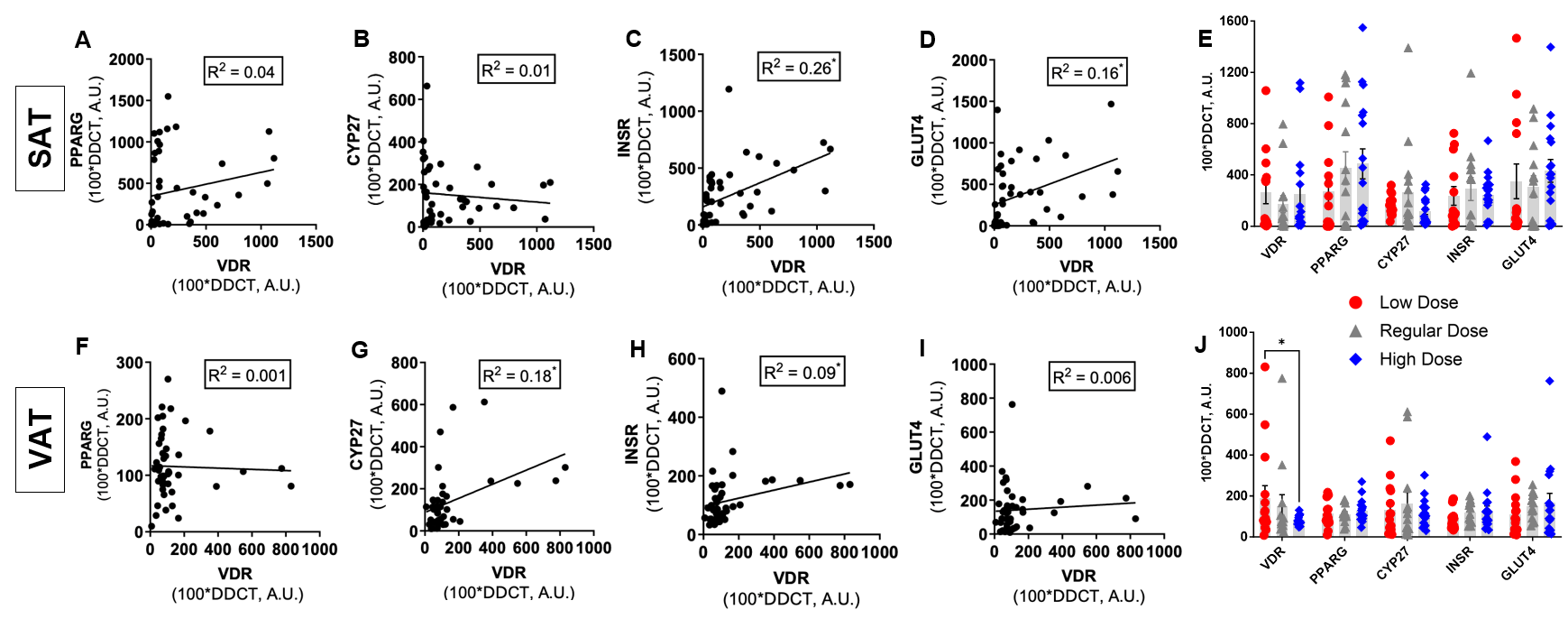

**SUPPLEMENTARY FIGURE 2:** VDR Gene expression correlates to glucose metabolism-related gene expression in adipose tissues. Subcutaneous Adipose Tissue (SAT): **(A)** *Vdr* vs *Pparg* gene expression, **(B)** *Vdr* vs *Cyp27a1* gene expression, **(C)** *Vdr* vs *Insr* gene expression, and **(D)** *Vdr* vs *Glut4* gene expression (calculated relative to the housekeeping genes *Gapdh* using the ΔΔCT method). **(E)** All examined SAT genes stratified by diet using raw Cq expression values. Visceral Adipose Tissue (VAT): **(F)** *Vdr* vs *Pparg* gene expression, **(G)** *Vdr* vs *Cyp27a1* gene expression, **(H)** *Vdr* vs *Insr* gene expression, and **(I)** *Vdr* vs *Glut4* gene expression (calculated relative to the housekeeping genes *Gapdh* using the ΔΔCT method). **(J)** All examined VAT genes stratified by diet using raw Cq expression values. **p*<0.05.

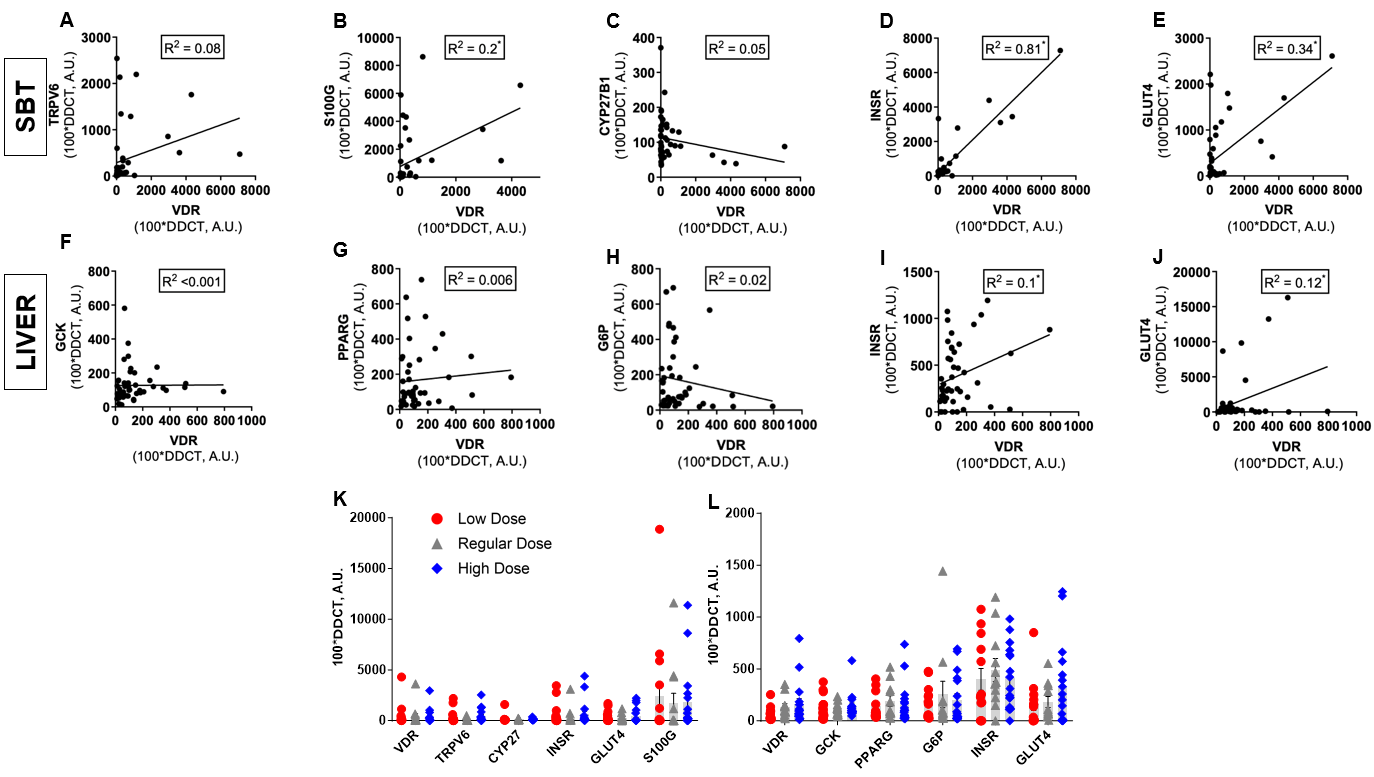

**SUPPLEMENTARY FIGURE 3:** VDR Gene expression correlates to glucose metabolism-related gene expression in organ tissues. Small bowel tissue (SBT): **(A)** *Vdr* vs *Trpv6* gene expression, **(B)** *Vdr* vs *S100g* gene expression, **(C)** *Vdr* vs *Cyp27b1* gene expression, and **(D)** *Vdr* vs *Insr* gene expression, and **(E)** *Vdr* vs *Glut4* gene expression (calculated relative to the housekeeping genes *Gapdh* using the ΔΔCT method). Liver: **(F)** *Vdr* vs *Trpv6* gene expression, **(G)** *Vdr* vs *S100g* gene expression, **(H)** *Vdr* vs *Cyp27b1* gene expression, and **(I)** *Vdr* vs *Insr* gene expression, and **(J)** *Vdr* vs *Glut4* gene expression (calculated relative to the housekeeping genes *Gapdh* using the ΔΔCT method). **(K)** All examined SBT genes and **(L)** liver genes stratified by diet using raw Cq expression values. **p*<0.05.
